## Supplemental Text for "A statistical framework for disease classification with scRNA-Seq Data"

### S1 Supplementary Text

#### S1.1 Library-size Normalization

To correct for differences in sequencing depth across samples before downstream analyses, we applied three commonly used library-size normalization strategies: Counts-Per-Million (CPM) [15], Upper Quartile (UQ) [16], and Median Ratio (MR) [17]. In all cases, normalization was performed by computing cell-type-specific size factors, allowing each cell type to be scaled independently and thereby accounting for heterogeneity in expression distributions across cell types.

- **Counts-Per-Million (CPM):** Let $C_{igl}$ denote the raw aggregated count for gene $g$ in cell type $l$ for sample $i$. The CPM-normalized value is $x_{igl} = 10^6 C_{igl} / \sum_{g,l} C_{igl}$.
- **Upper Quartile (UQ):** UQ normalization rescales each sample by its upper-quartile expression level. For sample $i$, we first compute the 75th percentile of its pseudobulk counts. If this upper-quartile value was zero (indicating many zeros due to dropout), the scaling factor was replaced by the median of the nonzero counts. The UQ-normalized value is then calculated as $x_{igl} = \frac{C_{igl}}{\mathrm{UQ}_i}$, where $\mathrm{UQ}_i$ is the adjusted sample-specific upper-quartile estimate.
- **Median Ratio (MR):** For each cell type $l$, genes with zero counts in any sample were excluded from calculation to avoid undefined geometric means. For the remaining genes, a reference expression value is first constructed as the gene-wise geometric mean across samples: $G_{gl} = \exp(\frac{1}{n} \sum_{i=1}^{n} \log C_{igl})$. For each sample $i$, the MR scaling factor is then computed as the median of the ratios $s_{il} = \mathrm{median}_g(\frac{C_{igl}}{G_{gl}})$, and normalized counts are obtained as $x_{igl} = C_{igl}/s_{il}$. Only genes with finite geometric means and nonzero counts across all samples contributed to the scaling factor.

#### S1.2 Dataset Overview

Dataset 1 : Perez et al. [20] present a initiative utilizing multiplexed single-cell RNA sequencing to systematically characterize the cellular and molecular landscape of systemic lupus erythematosus (SLE), a complex autoimmune disease with significant sex and ancestry disparities. The dataset contains over 1.2 million peripheral blood mononuclear cells (PBMCs) from 205 SLE cases and 131 healthy controls, providing one of the most comprehensive single-cell references for SLE to date. The study reveals that SLE pathogenesis is driven by coordinated transcriptional alterations across multiple immune cell types, with classical monocytes exhibiting most consistent and the highest levels of both pan-cell type and myeloid-specific type 1 interferon-stimulated genes (ISGs). Importantly, increased ISG signaling is a hallmark of SLE, higher ISG expression correlates positively with higher SLE Disease Activity Index (SLEDAI), and since classical monocytes are the main contributors to this signature, classical monocytes is a useful cell type for SLE prediction. Furthermore, naive CD4+ T cells are depleted from circulation in SLE, while cytotoxic CD8+ T cells show a significant clonal expansion in SLE.

Dataset 2 : Stephenson et al. [21] present a comprehensive single-cell multi-omics analysis that systematically characterizes the immune response landscape of COVID-19. We used a subset of the dataset, which contains 143 samples from sites Cambridge, Ncl, and Sanger. The study reveals that COVID-19 pathogenesis involves coordinated changes across multiple specific immune cell types, where classical CD14+ monocytes exhibit the strongest interferon response signatures and are significantly expanded in severe disease. CD16+ monocytes show elevated expression of complement gene transcripts (e.g., CD1QA/B/C), consistent with their proposed role in macrophage replenishment and inflammatory tissue remodeling in the lung. Cytotoxic CD8+ T cells also demonstrate increased clonal expansion and effector functions with COVID disease severity. Increased interferon signaling is a hallmark of COVID-19, with higher interferon response scores correlating positively with disease severity across all major immune cell populations, and since classical monocytes and myeloid cells are the main contributors to this signature alongside altered hematopoiesis marked by megakaryocyte-primed CD34+ stem cells, monocytes and myeloid populations are useful cell types for COVID-19 severity prediction and understanding disease pathogenesis.

Dataset 3: Pelka et al. [22] presents a comprehensive initiative utilizing single-cell RNA sequencing to systematically characterize the cellular and molecular landscape of colorectal cancer (CRC), a heterogeneous malignancy with striking differences in immune responsiveness between mismatch repair-deficient (MMRd) and mismatch repair-proficient (MMRp) subtypes. The dataset contains contains tumor and adjacent normal colon tissue samples from 62 patients (34 MMRd and 28 MMRp), with 36 patients contributing both tumor and adjacent normal tissue samples. The study reveals that CRC pathogenesis involves coordinated changes across multiple immune cell populations, with T cells and myeloid cells showing the most dramatic differences between tumor subtypes. CXCL13+ T cells are significantly enriched in MMRd tumors, while IL17+ T cells are preferentially found in MMRp tumors. Classical monocytes and macrophages demonstrate extensive reprogramming in tumors, expressing higher levels of inflammatory programs including glycolysis, alarmins, and neutrophil-recruiting chemokines, with these activation signatures being more pronounced in MMRd tumors. The identification of co-varying T cell and myeloid gene expression programs emerges as a hallmark of coordinated immune responses in CRC, with higher activities of CXCL13 T cell programs and myeloid activation signatures correlating with MMRd status and potential immunotherapy responsiveness, positioning T cell-myeloid interactions as critical determinants of tumor immunity and therapeutic susceptibility in colorectal cancer.
